# A Diabetic Mice Model For Studying Skin Wound Healing

**DOI:** 10.1101/2022.05.28.493835

**Authors:** Carlos Poblete Jara, Guilherme Nogueira, Joseane Morari, Thaís Paulino do Prado, Renan de Medeiros Bezerra, Bruna Bombassaro, Lício A. Velloso, William Velander, Eliana Pereira de Araújo

**Affiliations:** Faculty of Nursing; Faculty of Medical Sciences; Laboratory of Cell Signaling, Obesity and Comorbidities Research Center; University of Campinas, Brazil; Department of Chemical and Biomolecular Engineering, University of Nebraska-Lincoln, Lincoln, NE, USA

**Author notes:** Corresponding author: Eliana P. Araújo, Faculty of Nursing, University of Campinas, Brazil. Financial support: Coordination of Improvement of Higher-Level Personnel of Brazil (CAPES), FAPESP and Nebraska XXXX.

**Keywords:** animal model, high-fat-diet, streptozotocin, aging, wound healing

## Abstract

Advances in wound treatment depend on the availability of suitable animal models. All animal models try to reflect human wound healing problems. For acute wounds, it is easier to obtain adequate animal models, however, for chronic wounds such as those found in individuals with diabetic foot ulcer, approximations of the clinical picture become a challenge. Nowadays, the key points of wound healing processes are better understood, and therefore, therapeutic strategies can be developed to manipulate wound repair. Research efforts involves the development of therapies to aid in the treatment of impaired wound healing and, to improving normal wound healing to drive a process close to regenerative. To achieve a better animal model that is more appropriate for studying wound healing, six-week-old male C57BL/6 mice were separated into groups fed a Chow and High-Fat Diet for 0.5, 3, and 6 months, when part of the animals were induced to diabetes by streptozotocin. Then, mice were submitted to metabolic, molecular, and morphological analyses. We show that this model results in a severe metabolic phenotype with insulin resistance, reduced insulin expression, and glucose intolerance associated with obesity and, more importantly, skin changes. Furthermore, the skin phenotype, both structurally and transcriptionally, overlapped with conditions found in elderly patients with DM that reproduce the phenotype of most patients who develop diabetic foot ulcers.

## INTRODUCTION

Diabetes mellitus (DM) is a chronic metabolic disease characterized by hyperglycemia that results from deficiencies in insulin secretion and/or action. It is one of the major global health problems affecting over 463 million people worldwide and projecting an increase to 578 million by the end of 2023(1). As a chronic and refractory disease, DM affects every tissue and organ in the body, including the skin. Studies show that up to two-thirds of diabetic patients have skin problems at some point during their lifetime (2). There are several mechanisms behind DM-associated skin abnormalities, which include, but are not restricted to, oxidative stress, abnormal regulation of inflammatory products, impaired angiogenesis, and impaired growth factor production (3).

During the course of DM, two features seem central to explain the greater risk for development of skin disease: i, an accelerated skin aging (2, 4) and ii, an increased risk for development of secondary infections, which is particularly important in diabetic foot ulcers (DFU). DFU is the most common complication affecting DM patients; it increases the risk for development of osteomyelitis, which can lead to lower extremity amputations (5, 6) . Treating DFU has proven difficult as it results from a complex pathophysiology (6). Moreover, due to ethical concerns, trials for new therapeutic interventions in humans are limited, which delays the development of effective strategies to treat this condition. Therefore, testing new approaches to treat DFU must rely, at least during early phases, on the existence of appropriate experimental models.

Over the years, several experimental models have been used in studies aimed at evaluating interventions for treating DFU. These include spontaneous autoimmune DM in rodents, genetically induced DM models; diet-induced DM models and pharmacologically induced DM models (7–9). Regardless of the benefits obtained with each of these models, they all have particular limitations. One important limitation that occurs in virtually all models employed to date relies on the fact that they do not exhibit simultaneously two important features that are common to most patients with DFU; severe metabolic dysregulation and accelerated skin aging. Ideally, an experimental model for studying DFU should have close similarities to the clinical and pathological landscape of human DFU. In this study, we describe a model in which DM and aging result in skin alterations that display great similarities with those found in aging humans.

## RESULTS

### Diabetes induction

5-month old C57BL/6J mice were fed on HFD for 3 months when five low-dose injections of streptozotocin (STZ) (50 mg/kg, i.p.) were performed (Fig. 1a). Our results showed that after four weeks from STZ injections, HFD *diabetic* (HFD DM+) mice increased blood glucose levels up to 500mg/dL (489.7 mg/dL HFD DM+ vs 143 mg/dL age-matched non-diabetic mice, *p value* <0.0001). In addition, our result showed Chow diet did not affect fasting glycemia during the 8-month of analysis (Figure 1b, black bars). However, the 8-month old mice fed on HFD increased fasting glycemia compared to diet-paired 2.5-month old group (from 166 mg/dL in young to 206 mg/dL in older mice, *p value* 0.0268) (Figure 1b, blue bars). Moreover, the 8-month old HFD DM+ group increased even more fasting glycemia (3 fold more after 3 months streptozotocin injections) compared to diet-paired 2.5-month old counterparts (from 166 mg/dL younger to 490 mg/dL in older mice, *p value* <0.0001) (Figure 1b, red bars).

**Figure 1a.**
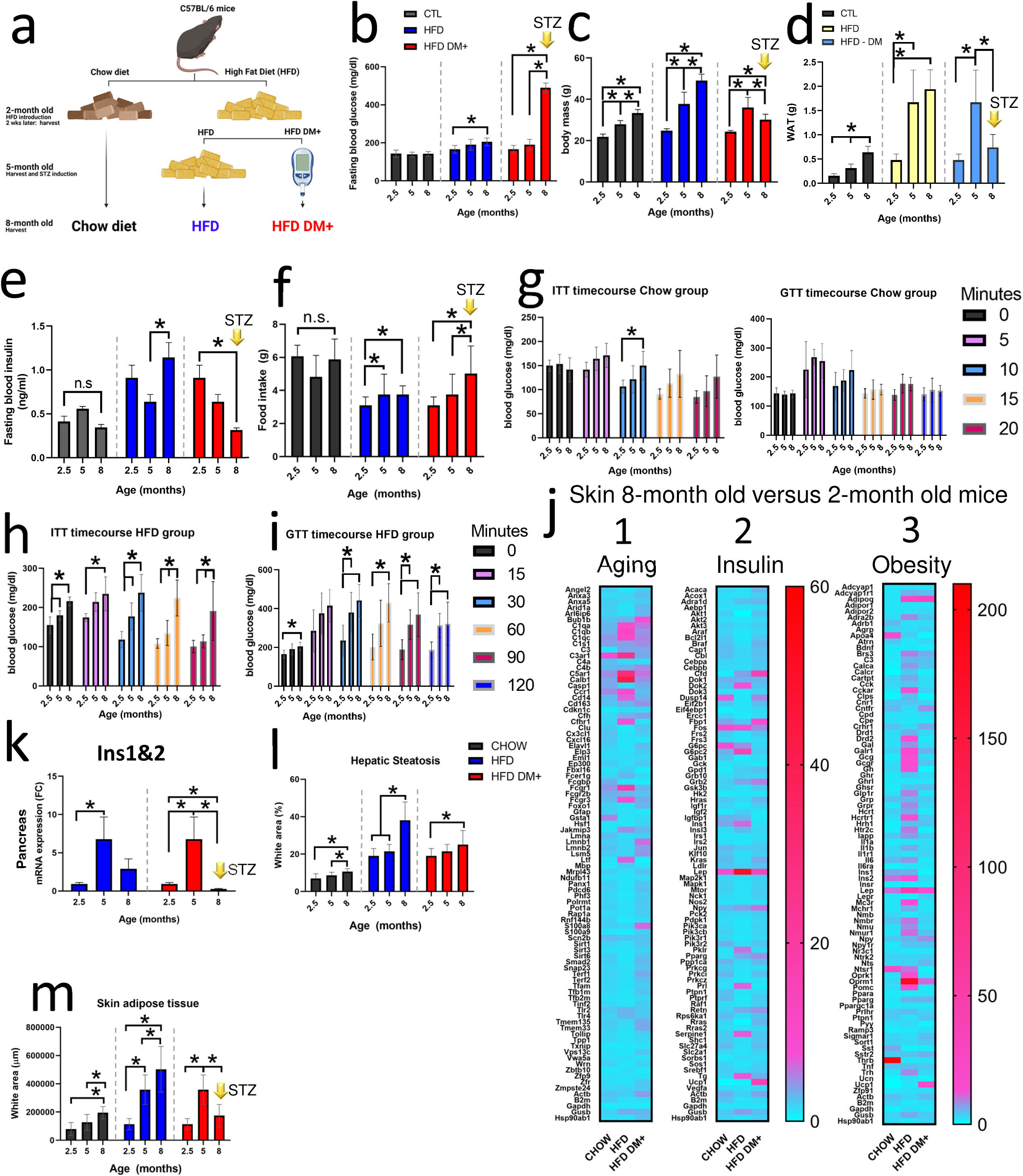
Schematic representation of experimental design. C57BL/6 mice were fed either with Chow Diet or High Fat diet. Some 5-month old HFD animals were treated with streptozotocin for Diabetes induction (HFD DM+ group) generating 2 HFD groups: a) HFD DM+ (T2DM-like) and b) HFD non-diabetic.

### Aging increases body mass and white adipose tissue deposition in mice

Aging affects body mass (Figure 1c). Independently of food characteristics, mice fed on Chow or HFD increased Body mass during the 8 months of analysis (Figure 1c and Table 1, *p value* <0.0001). 8-month old mice fed on Chow diet increased body mass (from 22g to 33g, *p value* <0.0001) at rate 60-80mg/day (Supplementary 1b). Moreover, 8-month old mice fed on HFD increased body mass (from 25g to 49g, *p value* <0.0001) at rate 130-170mg/day (Supplementary 1b). However, after streptozotocin injections, HFD DM+ mice decreased body mass (from 36g to 30g, *p value* 0.0173) at a rate 70 mg/day (Supplementary 1b).

**Table 1.**
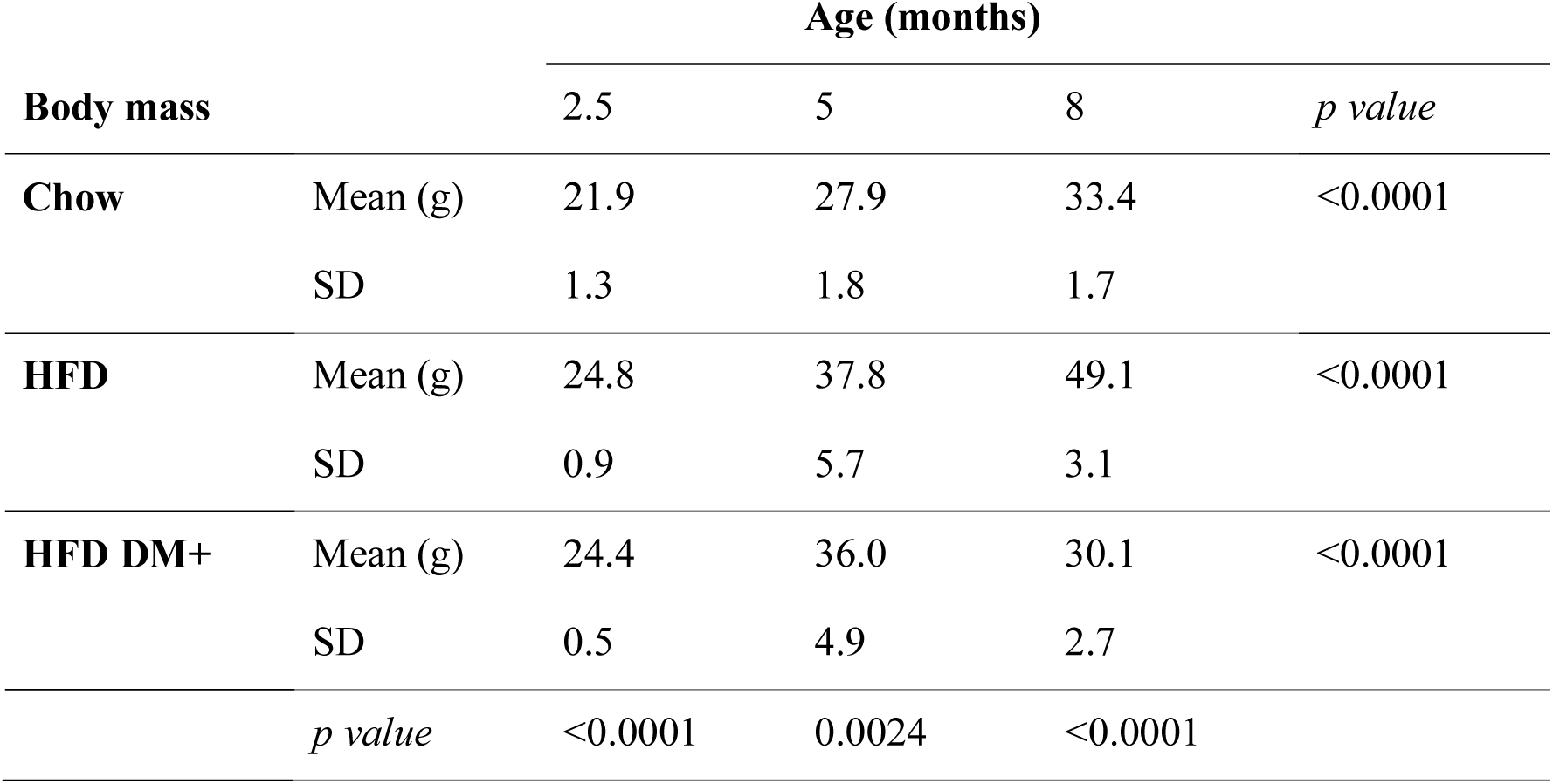

In addition, we found increased epididymal white adipose tissue (WAT) mass in older 5- and 8-month old mice fed with Chow diet compared to younger 2.5-month old mice fed on Chow (from 0.16g to 0.64g, *p value* <0.0001). This increase represents a 4-fold WAT mass in the 8-month old group fed with Chow diet compared with their younger counterpart (Figure 1d, black bars). We found this WAT mass increase following the Body Mass increase pattern (Supplementary 1a). Additionally, mice fed on HFD increased even more WAT mass (Figure 1d, blue bars). We found increased WAT in older 8-month old animals fed with HFD compared with younger HFD 2.5-month and 5-month old groups (from 0.48g to 1.94g, *p value* 0.0001). In the same way that Chow fed animals, the increased WAT mass represents 4-fold the WAT mass of younger mice fed on HFD (Figure 1d). As well as the Chow group, we found this increase in WAT mass also follows the Body Mass increment pattern (Supplementary 1a).

However, the biggest differences were found in older 8-month old HFD DM+. HFD DM+ mice decreased WAT mass compared to younger 5-month old diabetic mice (from 1.7g to 0.7g, *p value* 0.0005) representing 2.3-fold less WAT mass (Figure 1d, red bars). Interestingly, over the course of 3 months with induced diabetes beginning at 5 months’ age, this decrease in WAT mass also follows the Body Mass decline pattern of the HFD DM+ animals (Supplementary 1a).

### The hyperglycemic phenotype includes both insulin deficiency and polyphagia in HFD DM+

We explored how aging affects insulin levels and polyphagia. In 2.5-month old mice fed on chow diet showed 0.4 ng/mL insulin levels while 2.5-month old HFD mice showed 0.9 ng/mL insulin levels with no polyphagia. In 8-month old mice fed on chow diet showed 0.35 ng/mL insulin levels while 8-month old mice fed on HFD presented as 1.1 ng/mL insulin levels with no polyphagia. However, 8-month HFD-diabetic mice (HFD DM+) presented 0.3ng/mL insulin levels (Figure 1e) with polyphagia (Figure 1f).

8-month old HFD mice increased Food intake compared to 2.5-month old HFD mice (from 3g/day to 4g/day, *p value* 0.0232, Figure 1f). 8-month old HFD DM+ mice further increased Food intake to 5g/day (*p value* <0.0001, Figure 1f). In contrast, 8-month old Chow diet mice showed no differences in Food intake (Figure 1f).

### Aging impacts Insulin and Glucose tolerance in mice fed on HFD diet

We investigated the contribution of aging to insulin sensitivity by intraperitoneal injection of insulin (ITT) and glucose metabolism by intraperitoneal injection of glucose (GTT) in Chow, HFD, and HFD DM+ mice. Our results showed aging has a progressive and detrimental effect on insulin sensitivity and glucose metabolism in eight-month old HFD mice (Figure 1h-i). Increased glucose levels at 15, 30, 60, 90 and 120 min post insulin injection occurred compared to 2.5-month old HFD mice (*p value* <0.05, Figure 1h). GTT results in 8-month HFD mice also showed increased glucose levels 30 min after glucose challenge compared to 2.5-month HFD mice (*p value* <0.05, Figure 1i). Our results in 2.5, 5 and 8-month Chow fed mice showed no differences in the clearance pattern of plasma glucose except after 10 min post insulin injection (from 106 mg/dL to 150 mg/dL, *p value* 0.0260, Figure 1g left).

### Aging impacts gene expression associated with insulin pathway, obesity and aging

To better understand the effect of aging in the skin of Chow, HFD and HFD DM+ mice, we screened three different gene arrays: Aging, Insulin Pathway and Obesity.

**Column 1**. Figure 1J shows the expression of age-associated genes in the skin of 8- month old animals relative to 2-month old counterparts. 8-month old Chow animals overexpressed *C3ar1* (8-fold), *Gsta1* (5-fold), and *Ccr1* (4-fold). In addition, 8-month old HFD animals specifically overexpressed *Calb1* (42-fold), *C5ar1* (30-fold), and *C3ar1* (26- fold). Skin from 8-month old HFD DM+ mice specifically overexpressed *Bub1b* (7-fold), *S100a8* (6-fold), and *Lmnb1* (5-fold).

**Column 2**. Figure 1J shows the expression of Insulin-associated genes in the skin of 8-month old animals relative to 2-month old counterparts. 8-month old Chow animals overexpressed *Lep* (9-fold), *Fos* (9-fold), and *G6pc* (8-fold), the HFD animals overexpressed *Lep* (50-fold), *G6pc2* (15-fold), and *Fos* (7-fold), and the HFD DM+ specifically overexpressed *Bub1b* (7-fold), *S100a8* (6-fold), and *Lmnb1* (5-fold).

**Column 3**. Figure 1J shows the expression of Obesity-associated genes in the skin of 8-month old animals relative to 2-month old counterparts. 8-month old Chow animals specifically overexpressed *Thrb* (207-fold), *Ntsr1* (20-fold), and *Apoa4* (16-fold), the HFD animals overexpressed *Oprm1* (182-fold), *Lep* (48-fold), and *Adipoq* (25-fold), and the HFD DM+ overexpressed *Adipoq* (28-fold), *Ucp1* (15-fold), and *Lep* (13-fold).

### Aging affects pancreatic Insulin 1 and 2 expressions in mice fed on HFD

*Ins1* and *Ins2* genes encode for insulin 1 and 2, peptides that are vital in the regulation of carbohydrate and lipid metabolism. Our result showed that total *Ins1&2* gene expression increased in **pancreatic** tissue in 5-month old mice fed on HFD (Figure 1k). After streptozotocin injections, the HFD DM+ mice decreased pancreatic total *Ins1&2* gene expression (Figure 1k).

### Aging affects the accumulation of intrahepatic and dermal fat in HFD and Chow fed mice

Both the accumulation of intrahepatic fat (Hepatic steatosis) and insulin resistance are associated with liver metabolic dysfunction (10, 11). For this reason, we explored intrahepatic fat percentage in HFD DM+. Figure 1l shows the histological percentage of intrahepatic fat. After 8 months on Chow diet, intrahepatic fat increased from 7% to 11% when comparing to 2-month old mice (*p value* <0.0001). 8-month old HFD mice had increased intrahepatic fat percentage from 19% to 38% (*p value* <0.0001). HFD DM+ animals increased from 19% to 25% (*p value* 0.0148).

Dermal white adipose tissue (dWAT), occurs in the dermis underlying the reticular dermis (12), and participates in thermogenesis, wound healing and immune defense against infection (13). We investigated dermal fat deposition in mice fed on Chow, HFD and HFD DM+.

Figure 1m shows the histological presence of dWAT. 8 months Chow and HFD animals increased dWAT relative to both 2- and 5-month old mice (*p value* <0.0001). In contrast, HFD DM+ mice decreased dWAT relative to 5-month old HFD mice (*p value* <0.0001).

### HFD and hyperglycemia affects body mass in HFD DM+ mice

In our previous studies of mice less than 5-month age, we have shown that HFD affects body mass (14–17) (). Figure 2a shows that after only 2 weeks on HFD, mice had 13% increased body mass as compared to the Chow diet group (24.6g HFD vs 21.9g Chow, *p value* <0.0001). 3 months additional feeding on HFD resulted an increase of 35% of body mass compared to Chow diet (37.8g HFD vs 27.9g Chow, *p value* 0.0024). A total of 6 months HFD increased body mass by 47% compared to Chow diet mice (49.1g vs 33.5g, *p value* <0.0001). In contrast, HFD DM+ after a total of 6 months on HFD had decreased 39% body mass compared with their non-diabetic HFD counterparts (30.1g HFD DM+ vs 49.1g HFD, *p value* <0.0001).

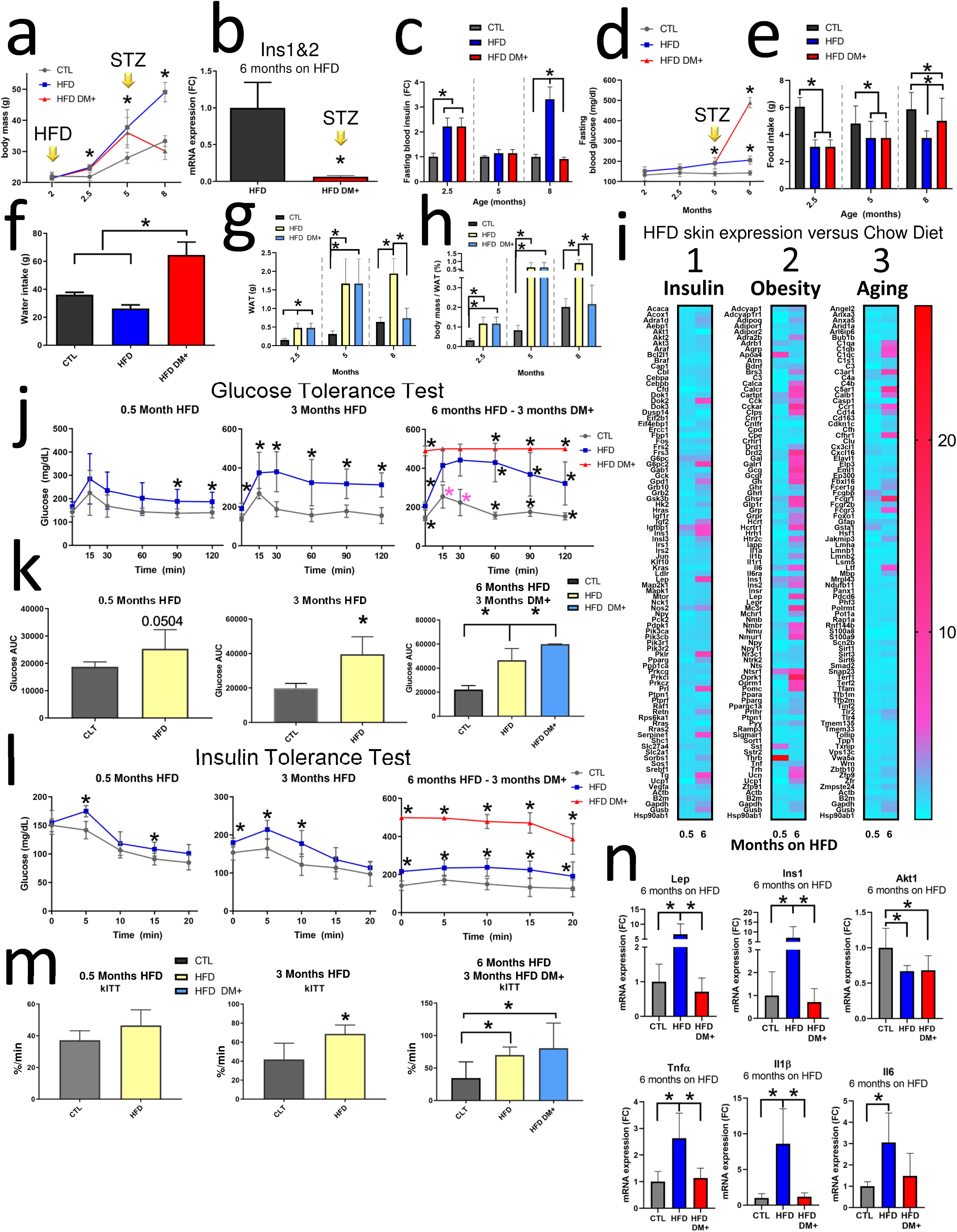

### HFD DM+ have both decreased insulin expression and circulating insulin

Our result showed that after post Streptozotocin treatment, HFD DM+ decreased Pancreatic *Ins1* and *Ins2* expression (0.05-fold change, *p value* 0.0335) at 8 months of age (Figure 2b). Figure 2c shows fasting blood insulin levels. Mice fed for only 0.5 months on HFD beginning at 2 months age had increased fasting blood insulin levels compared to Chow animals (0.41ng/mL vs 1.3ng/mL, *p value* 0.0374). Mice after a total of 6 months on HFD increased insulin levels to 1.1ng/mL while Chow animals had 0.3 ng/mL (*p value* 0.0017). 8-month HFD DM+ (6 months on HFD) showed 0.3ng/mL.

We found mice fed on HFD for 3 months increased 37% Fasting blood glucose compared to Chow diet mice (190mg/dL vs 139mg/dL, *p value* 0.0021) (Figure 2d). After 6 months on HFD, mice had a 41% increased glucose levels compared to Chow mice (201mg/dL vs 142mg/dL, *p value* <0.0001). In contrast, HFD DM+ had 144% and 243% increased Fasting blood glucose when compared with age and HFD- and Chow, respectively (490mg/dL vs 201mg/dL vs 142mg/dL, *p value* <0.0001).

### Diabetic mice showed polyphagic food intake and polydipsic water intake

Our results showed HFD DM+ mice increased daily Food intake (Figure 2e). HFD DM+ mice displayed a 25% increased daily Food intake compared with their non-diabetic HFD counterparts (5g vs 4g, *p value* 0.0021). We found no differences in water intake (9mL vs 6mL, *p value* 0.0718) between Chow diet and HFD mice (Figure 2f). However, HFD DM+ showed *polydipsic* water intake compared either to Chow diet or their non-diabetic HFD counterparts (16g vs 9g, *p value* <0.0001) (Figure 2f). This difference in diabetic mice represents a 44% more daily water intake compared to the HFD mice and 63% more water intake compared to mice fed on Chow diet.

### HFD DM+ have decreased epididymal White Adipose

HFD mice increased WAT mass compared to Chow diet (480mg vs 156mg, *p value* 0.0002) (Figure 2g). This difference represents a 207% more WAT mass in HFD animals compared to the Chow group. An additional 3 months on HFD resulted a further 431% increase of WAT (1674mg vs 315mg, *p value* 0.0007) compared to Chow diet (Figure 2g). After 6 months on HFD, mice further increased WAT mass by 205% (1942mg vs 638mg, *p value* <0.0001) compared to Chow diet (Figure 2g). In contrast, 8-month HFD DM+ mice decreased WAT mass by 161% compared to their non-diabetic HFD counterparts (743mg vs 1942mg, *p value* <0.0001) (Figure 2g). We identified similar results of White adipose tissue mass after Body Mass correction (Figure 2h).

### HFD and HFD DM+ mice present chronic glucose intolerance

GTT results showed increased plasma glucose levels at 90 min (138mg/dL vs 189mg/dL) and 120 minutes (140mg/d vs 187mg/dL) after 0.5 months of HFD compared with Chow diet mice (*p value* <0.05) (Figure 2j). After 3 months of HFD feeding, mice increased plasma glucose levels at each time point of GTT sampling (0, 15, 30, 60, 90 and 120 min) (*p value* <0.05). 8-month old HFD mice (HFD for 6 months) also had increased plasma glucose levels at all time of GTT sampling (*p value* <0.0001). HFD DM+ mice presented higher plasma glucose levels (498mg/dL vs 364mg/dL) at all GTT time points compared to their age-matched HFD group (*p value* <0.0001). There were no differences in the AUC (from plasma glucose levels) of animals fed on HFD for 0.5 months (Figure 2k). After 3 months on HFD, the AUC increased 99% compared to Chow animals (*p value* 0.0010) (Figure 2k). HFD DM+ mice also showed a 170% AUC increase relative to the Chow diet group (*p value* <0.0001)(Figure 2k).

### HFD and HFD DM+ mice present chronic Insulin resistance

To determine the whole-body sensitivity to insulin, we measured blood glucose levels after intraperitoneal insulin administration (Figure l-m). ITT results showed that after 0.5 months of HFD plasma glucose levels increased at 5 min (175mg/dL vs 142mg/dL) and 15 min (108mg/dL vs 91mg/dL) compared to mice fed on Chow diet (*p value* <0.05) (Figure l). After 3 months on HFD the plasma glucose levels increased at 0 min (180mg/dL vs 153mg/dL), 5 min (214mg/dL vs 164mg/dL), and also 10 min (177mg/dL vs 122mg/dL) (*p value* <0.05).

After 6 months on HFD, the plasma glucose levels were increased at all-time points (*p value* 0.0001). 8-month old HFD DM+ mice (6 months on HFD) also presented higher plasma glucose levels at all-time points (*p value* <0.0001). A constant rate of glucose disappearance was observed (kITT) (Figure 2m) where higher values indicate greater tissue insulin resistance (18, 19). As results, no kITT differences were observed after 0.5 months of HFD feeding (*p value* 0.0783) (Figure 2m) but kITT increased after 3 months on HFD (*p value* 0.0071) (Figure 2m). This kITT difference represents a 41% more in insulin resistance in HFD animals compared to the Chow group. HFD DM+ showed increased kITT values compared to age-matched Chow diet groups (*p value* 0.0274), but not when compared to the HFD group (Figure 2m). This kITT difference in HFD DM+ mice represents a 133% increase in insulin resistance compared to Chow diet mice but no difference with the HFD group (p valor 0.7867) (Figure 2m).

### Diet and glycemia impact gene expression associated with insulin pathway, obesity and aging

**Column 1.** Figure 2i shows the expression of insulin-associated genes in the skin of 8-month old HFD animals relative to 8-month old Chow counterparts. After 2 weeks on HFD, skin tissue increased *Igfbp1* (3.2-fold), *G6pc* (2.5-fold), and *Ins1* (2-fold) more transcripts than their age-matched Chow diet counterparts. After 6 months on HFD, skin tissue overexpressed *G6pc2* (8.7-fold), *Lep* (8.5-fold), and *Ins1* (6 fold) more transcripts as well as downregulation of Akt1 (0.01 fold) when compared to their age-matched Chow diet counterparts (Figure 2i).

**Column 2.** Figure 2i shows the expression of obesity-associated genes in the skin of 8-month old HFD animals relative to 8-month old Chow counterparts. After 2 weeks on HFD, skin tissue increased *Thrb* (26-fold), Apoa4 (8.5-fold), and *Sst* (4-fold). After 6 months on HFD, skin tissue overexpressed *Oprk1* (17-fold), Drd2 (11-fold), and *Mc3r* (11-fold) (Figure 2i).

**Column 3.** Figure 2i shows the expression of aging-associated genes in the skin of 8- month old HFD animals relative to 8-month old Chow counterparts. After 2 weeks on HFD, only 3 of 89 genes were >2-fold expressed: *Fcgbp* (3-fold), *Gsta1* (2.8-fold), and *Fcgr1* (2.3- fold) had increased expression. After 6 months on HFD, skin tissue from HFD mice increased expression of *Fcgr1* (16-fold), *C5ar1* (11-fold), and *C3ar1* (9-fold) (Figure 2i).

To confirm the above array results from skin samples, we assessed individual gene expression by PCR. Our PCR results showed increased for HFD (after 6 months feeding) of the expression of *Lep* (5.6-fold) as compared to Chow diet mice (*p value* 0.0011) and also 5.9 fold more *Lep* expression compared to HFD DM+ (*p value* 0.0011). After 6 months on HFD, an increase of 11.2-fold of *Ins1* as compared to Chow diet mice (*p value* 0.0003), and 10-fold as compared to HFD DM+ (*p value* 0.0001). *Akt1* was downregulated in both HFD and HFD DM+ mice (*p value* 0.0291). The inflammatory gene markers after, mice increased *Tnf*[(*p value* 0.0008), *Il1*β (*p value* 0.0004) and *Il6* (*p value* 0.0156) after 6 months on HFD (Figure 2n).

### HFD and hyperglycemia affect intrahepatic fat deposition in mice

Our results showed that HFD and hyperglycemia both are associated with accumulation of intrahepatic fat percentage in mice (Figure 3a, c). Mice fed for 0.5 months on HFD had increased intrahepatic fat compared to age-matched Chow fed mice (19% vs 11%, *p value* <0.0001). Mice fed for 3 months on HFD also had increased intrahepatic fat compared to age-matched Chow fed mice (21% vs 13%, *p value* <0.0001). After 6 months on HFD, mice further increased intrahepatic fat compared to age-matched Chow fed mice (38% vs 9.3%, *p value* <0.0001). In contrast, HFD DM+ had decreased intrahepatic fat percentage as compared to age-matched HFD fed mice (38% vs 25%, <0.0001) (Figure 3a,c).

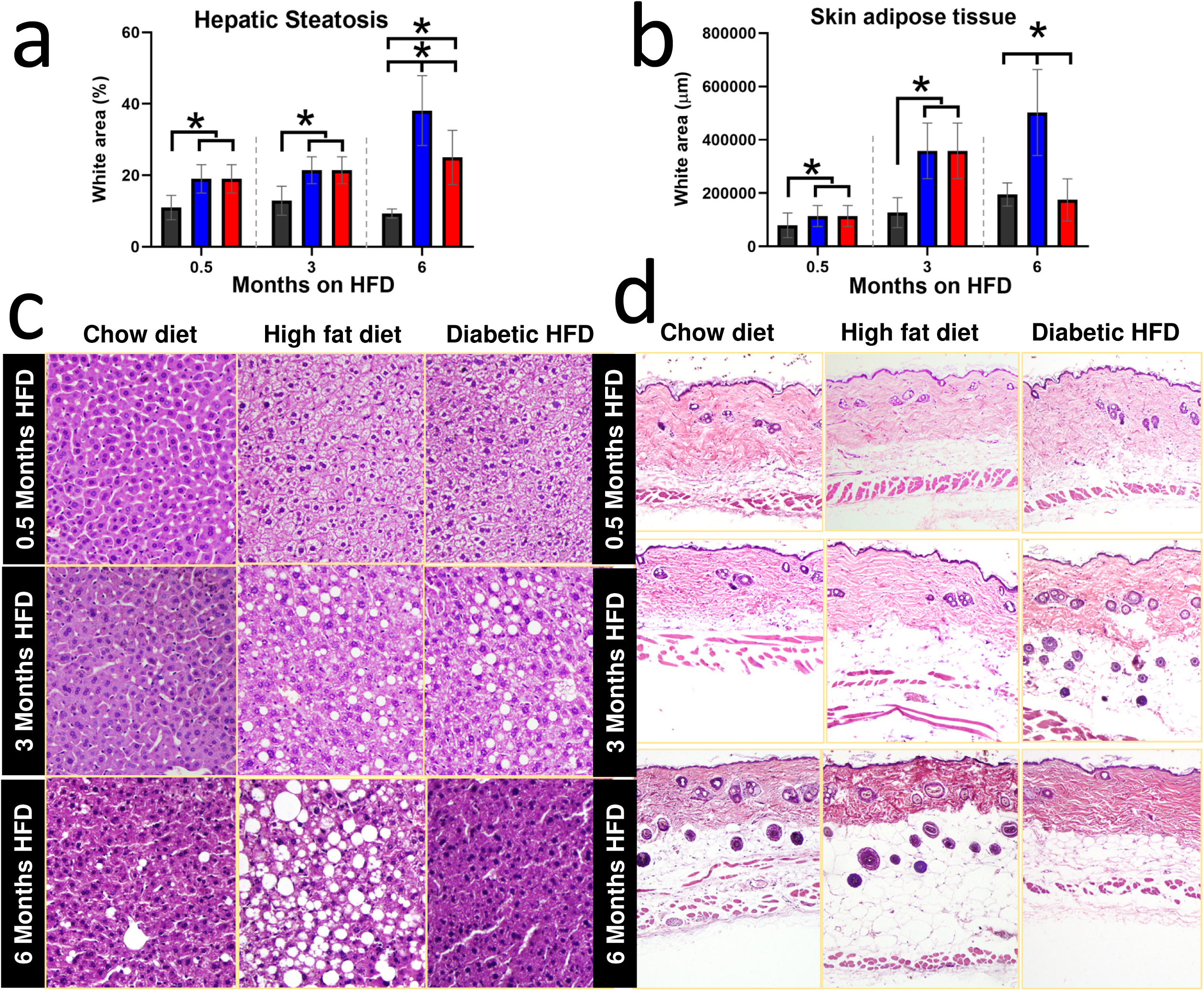

### HFD and hyperglycemia affect dermal fat deposition in mice

Our histological analysis showed that HFD and HFD DM+ had different levels of accumulated dWAT (Figure 3b,d). Both mice with 0.5 months and 3 months on HFD had increased dWAT as compared to the Chow fed mice (*p value* <0.05). In contrast, HFD DM+ mice had decreased dWAT as compared to HFD group (*p value* <0.0001, Figure 3b,d).

### A comparison of gene expression of diabetic human and HFD DM+ mice in skin

A Mouse Insulin Pathway array (Qiagen, PAMM-030Z) was used to compare 20-gene expression of skin samples with diabetic human and HFD DM+. Relative to non-diabetic skin samples, our analysis identified 10-common genes: *RRAS, SLC27A4, CFD, EIF4EBP1, GRB2, SORBS1, PTPN1, GSK3B, HRAS* and *BRAF* (Figure 4a).

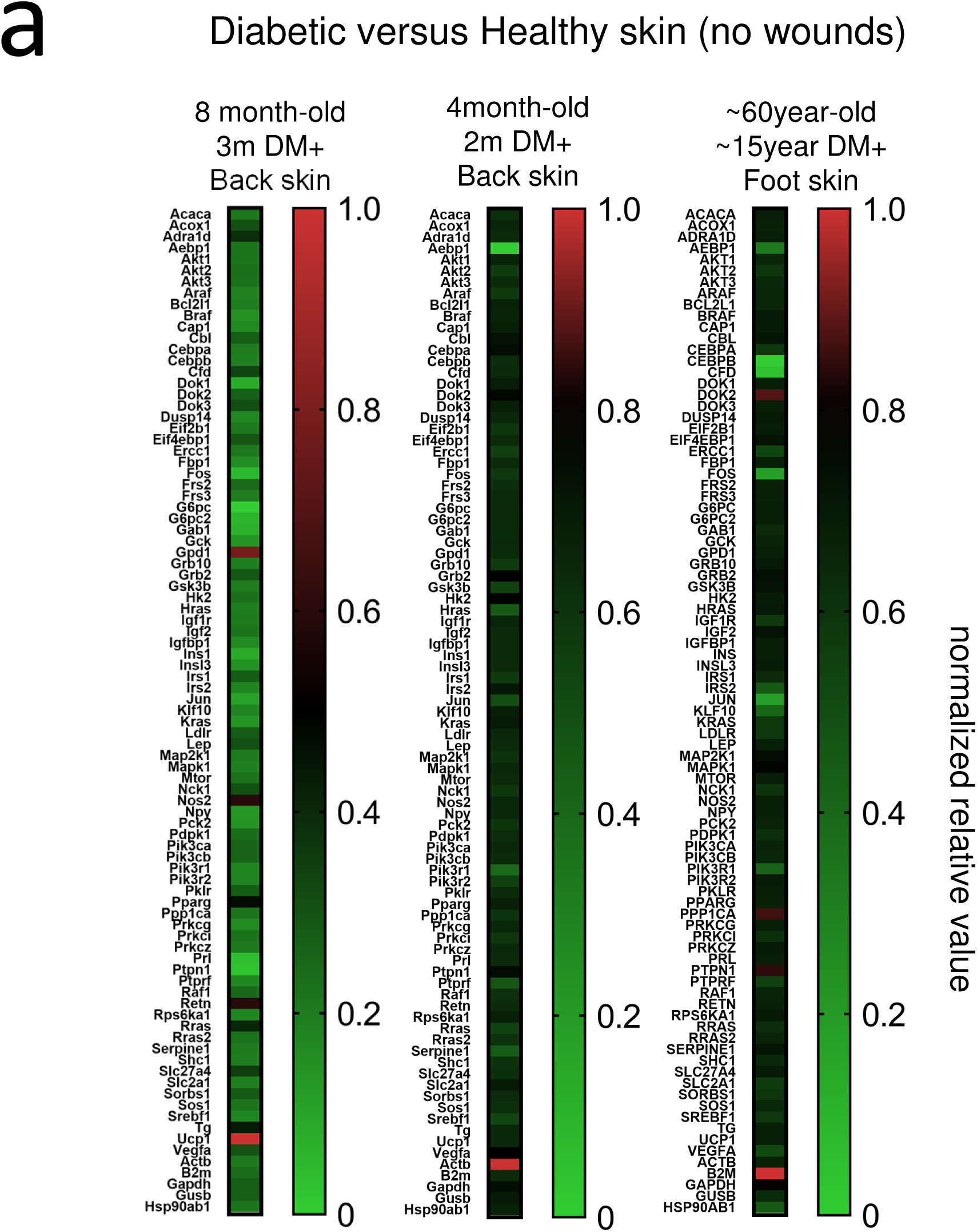

We compared this 8-month old HFD DM+ model data to previously published data of 4- month old STZ-induced diabetic non-HFD mice. We identified four common genes increased between older and younger STZ-induced diabetic mice: *Pparg*, *Dok3*, *Vega* and *Grb2* (Figure 4a).

### HFD DM+ showed delayed wound healing rate

We confirmed hyperglycemic status before performing wound experiment in 8-month old mice. HFD DM+ group presented higher fasting blood glucose compared to Chow or HFD mice either the same day of wounding or 12 days post wounding (p value <0.05, Figure 5a). HFD DM+ mice showed increased Wound area compared to Chow diet at 6 days (28mm^2^ vs 24mm^2^), 8 days (19mm^2^ vs 13mm^2^), and 10 days (11mm^2^ vs 8mm^2^) post-wounding (p value <0.05, Figure 5b,e,g). These differences in HFD DM+ mice represent respectively more wound area compared to Chow animals and suggested a slower healing process in the HFD DM+ group. Additionally, qualitative analysis suggested HFD and diabetic HFD increased cellularity in 12-day wounded skin tissue (Figure 5g). Microscopically, 8-months old Chow, HFD and HFD DM+ mice showed complete re-epithelialization at 12 days post wounding (Figure 5g).

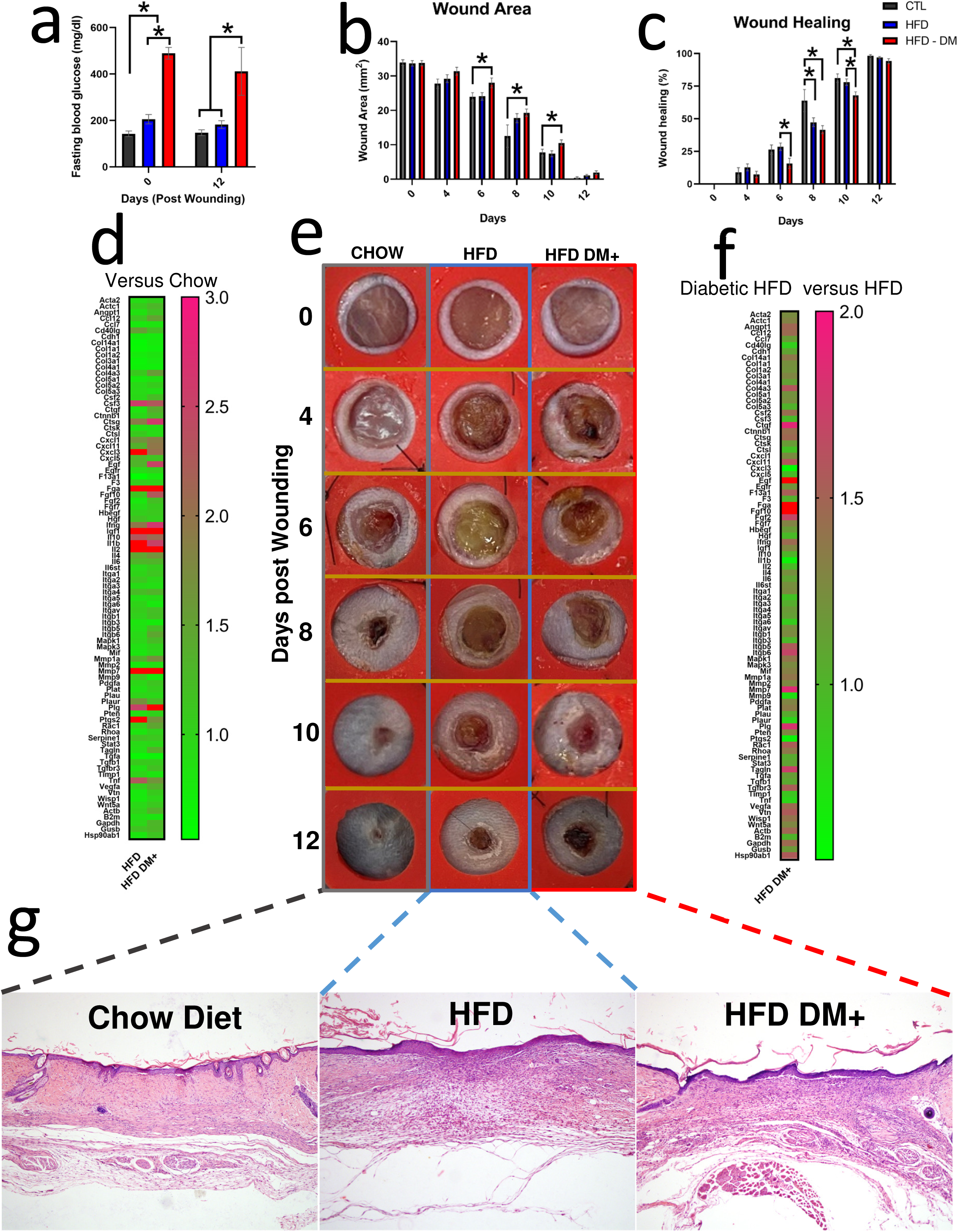

### Diet and glycemia affect gene expression of wound healing pathway

Figure 5d shows the expression of wound healing-associated genes in 12 days post wounding skin tissue of 8-month old HFD and HFD DM+ animals relative to 8-month old Chow counterparts. After 6 months on HFD, mice increased *Il1b, Ptgs2, Il10, Tnf, Mmp1,* and *Cxcl1* as well as decreased *Tgfa* and *Col14a1* as compared to age-matched Chow diet counterparts (p value <0.05). HFD DM+ skin increased Egf, Fgf10, Mmp1a, Ccl12 as well as downregulation of Col1a1 when compared to age-matched Chow diet counterparts (p value <0.05). Additionally, we explored HFD DM+ gene expression compared to HFD group as baseline. As a result, 12 days post wounding skin tissue increased *Fgf10, Egf, Ctgf, Itgb6*, and *Fgf2*.

To confirm the above array results from 12 days post wounded skin samples, we assessed individual gene expression by PCR. Figure 6a showed HFD DM+ increased *Ins1* and *Ins2* as well as decreased *Col1a1* gene expression (p value <0.05). HFD mice increased *Tnfa* and *Il1b* gene expression compared either to Chow or HFD DM+ groups (*p value* <0.05)

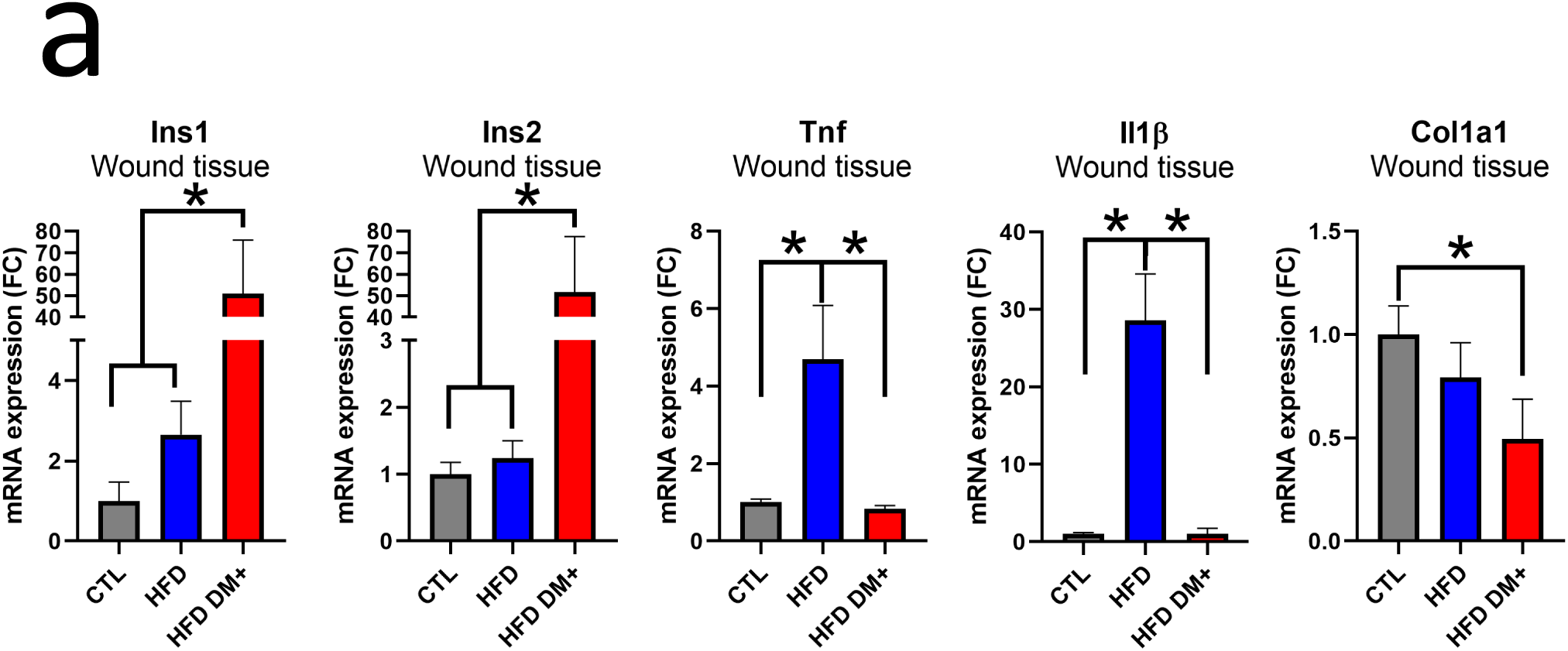

## DISCUSSION

Diabetic Foot Ulcers (DFU) are the most common skin condition associated with diabetes complications and these are accompanied with high risk of infections and eventually amputations (31–33). In this way, the development of new therapeutic interventions for promoting faster and more effective healing are needed. It is well know that testing new biotherapeutic is challenging with several ethical barriers. Thus, in early phases of testing, animal models provide a fast and effective approach to identify new pharmacological candidates. Most animal models used do not match clinical conditions frequently seen in humans with diabetes and DFU. In this way, our objective was to describe a mice model for skin wound healing assessment with diabetic and obese features, and then, offers a platform for accelerating DFU new treatment testing.

Diabetes Mellitus (DM) is a chronic metabolic disease associated with accelerated aging and resulting in the damage of several tissues and organs (22–24). For instance, skin of diabetic patients undergoes key alterations at metabolic, vascular, immunological, and neural levels, especially in poorly controlled DM individuals (25–27). However, the processes correlating these alterations are not yet entirely elucidated (4, 28). Our results suggest that HFD and hyperglycemia follows some of these key alterations at metabolic level, similar to poorly controlled DM patients. In long term DM, patients present both glucose intolerance and insulin resistance associated with insufficient capacity of pancreatic beta cells to produce insulin to cope with physiological demand (20, 21). We showed that an experimental model that associates diet-induced obesity, aging and pharmacological induction of diabetes mellitus results in a severe metabolic phenotype with insulin resistance, reduced b-cell insulin expression and glucose intolerance. We described the metabolic phenotype of an experimental model that can be used to study skin wound healing in aging and DM. We suggest that these pathological characteristics in our model are similar to key changes in human skin.

In our animal model, we observed important aging-associated transcripts changes that could be interpreted as alterations in the skin (Figure 4a). For example, we observed decreased in transcripts involved in insulin action and obesity. Additionally, we observed changes in pro-inflammatory cytokines that are crucial in regulating the inflammatory process during cutaneous wound healing. Our results showed abnormal levels of these cytokines suggesting a prolonged inflammatory phase and delaying the wound healing progression (6, 29, 30). Specifically, we showed increased *Tnf*α and *Il1*β gene expression in wounded and unwounded skin tissue. In addition, we found decreased *Col1a1* expression in wounded skin from diabetic mice. Interestingly, 8-month old mice increased *Ins1* expression after 6 months on HFD (Supplementary 1c).

After 12 days post wounding, transcriptomic, micro and macro wound evaluation suggest HFD DM+ mice are still resolving proliferative phase compared to HFD mice. Micro and Macroscopic evaluation showed increased wound area and delayed wound healing rate, altogether, suggesting a HFD pro-inflammatory phenotype and a healing-delayed HFD DM+ phenotype in hyperglycemic mice fed on HFD.

Usually mice models with severe diabetes, develop such condition early in life, thus, the important component of aging is missing (7–9). Previous models of DM induced by the consumption of energy dense diets showed chronic conditions that can be sustained until late life; however, metabolic abnormalities are generally mild (34–36). Here, we described a mouse model that enhances metabolic severity (pharmacological-induced DM) to the aging process occurring on a long-term HFD feeding. We hope that our model could be helpful for researcher as platform for accelerating DFU new treatment testing.

## CONCLUSION

Here we proposed a model in which mice were submitted to a protocol aimed at producing a severe DM features associated with obesity and aging. In this way, we described a phenotype of diabetic patients developing DFU. We showed that mice submitted to this intervention presented severe metabolic abnormalities associated with aging and hyperglycemia. Moreover, the skin phenotype, both structurally and transcriptionally overlapped with the conditions found in aging diabetic individuals. Thus, we have produced an animal model that could be used as platform for studying HFD and hyperglycemic-associated skin conditions.

## METHODS

### RT2 Profiler PCR arrays and Bulk expression

Gene expression was analyzed as previously described (37) . Briefly, Gene expression was obtained using four different 96-well RT2 Profiler^TM^ PCR Arrays, Mouse Wound Healing (No. PAMM-121Z), Mouse Insulin Pathway (PAMM-030ZC), Mouse Obesity (PAMM- 017ZC-12) array and Mouse Aging (PAMM-178ZC) from Qiagen (Maryland, USA), according to the manufacturer’s instructions. For each treatment group and time point, the array was run three times. RT-PCR was run on a StepOnePlus™ Real-Time PCR system (Thermo Fisher Scientific). For data normalization, *Hsp90ab1* and *Gapdh* average was selected as the reference gene based on data from the Mouse Housekeeping Genes RT² Profiler™ PCR Arrays and subsequent calculation of M values using the geNorm software (Biogazelle NV, Gent, Belgium). StepOne Software v.2.3 was used for data analyses, and gene expression was calculated with the ΔΔCt method. A gene was assumed to be differentially expressed if there was at least a twofold difference expression. For skin samples, we used *n* sample =2 pooling 2-3 animals per sample. For wound healing arrays, we used *n* samples =3 pooling 2-3 animals per sample. For pancreatic *Ins1&2* mRNA expression, we obtained an *Ins1* and *Ins2* mean expression using *Gapdh* and *Rpl0* mean as endogenous control. For dermal *Lep, Ins1, Akt1, Tnf*α*, Il1*β, and *Il6*, we used *Gapdh* as endogenous control. In Figure, 1j we used 2-month old skin samples as reference. In Figure, 2i we used skin from Chow diet mice as reference. We used public available single cell RNA sequencing datasets as “bulk” expression of diabetic skin from mice and human diabetic. Briefly, we retrieved gene expressions from diabetic samples (38, 39) (human and mice) according to the gene list of the Mouse Insulin Pathway array (PAMM-030ZC). For Fold change, we used non-diabetic “healthy” skin as reference (39, 40). As previously described (41), we used *In silico* analyses were performed using a DELL workstation with 16 GB RAM and four-cores Intel i7 processor. Sample expression matrices were downloaded from Gene Expression Omnibus Archive: GSE142471 and GSE165816. Cells were filtered by their total number of reads, by their number of detected genes and by their mitochondrial percentage. For mice, we used nFeature_RNA < 6,000, nCount_RNA < 40,000, percent.mt < 25% settings. For humans, we used nFeature_RNA < 2,500, nCount_RNA < 8,000 and percent.mt < 20% settings. Samples were processed in Seurat v3.1.5 using the default Seurat workflow (42). For clustering and visualization, we used the default Seurat pipeline and Prism Graphpad (8v) heatmaps.

### Experimental model

Six-week-old male C57BL/6 mice were obtained from the Animal Facility of the University of Campinas. Mice were kept in groups (5 mice per cage) at 21 ± 5 °C, in 12/12 h light/dark cycle, with water and chow *ad libitum*. Next, mice were kept in individual cage since 1 week before testing, measurement, and tissue harvest. In all experiments, control and intervention group mice were treated in the same experimental settings. All experiments were conducted according to the “Guide for the Care and Use of Laboratory Animals of the Institute of Laboratory Animal Resources, US National Academy of Sciences” and were approved by the Ethics Committee (CEUA IB/UNICAMP n° 5425-1/2019).

### Dietary interventions

Eight-week-old mice were separated into seven groups. Three groups fed with Chow diet (base-line control). The remaining four groups were fed with High-Fat diet (HFD) for 2, 12 or 24 weeks (diets composition in Table 2, experimental design in Figure 1a). Then, mice were subjected to lethal anesthesia and tissues specimens were extracted for analyses.

**Table 2.** Macronutrient component of the diets.

| **Components** | **Chow (g)** | **High-fat diet (g)** |
| --- | --- | --- |
| Starch | 427.5 | 199.5 |
| Protein (casein 85%) | 200 | 200 |
| Dextrinized corn starch | 132 | 132 |
| Sucrose | 100 | 100 |
| Soyabean oil | 40 | 40 |
| Lard | 0 | 228 |
| Fiber (celulose) | 50 | 50 |
| Mineral mix (ain-93) | 35 | 35 |
| Vitamin mix (ain-93) | 10 | 10 |
| L-cystine | 3 | 3 |
| Choline bitartarate | 2.5 | 2.5 |
| **TOTAL** | **1000** | **1000** |

### Diabetes Mellitus induction protocol

At 20 weeks of age (5-month old), one group on HFD was induced to diabetes mellitus (DM) with Streptozotocin (STZ). Mice were fasted for 4h before daily intraperitoneal injections of STZ (50 mg/kg). STZ was daily and freshly dissolved in 0.1 M sodium citrate buffer, pH 4.5, and the dose volumes for each mouse were calculated by *Labinsane* app (43) STZ injections were made for five consecutive days (low multiple-dose protocol) (44). HDF and Chow diet were injected sodium citrate buffer as control group.

Induced DM state was assessed after four weeks using an OptiumTM mini (Abbott Diabetes Care, Alameda, CA, USA) handheld glucometer with appropriate test strips. Blood glucose levels were measured by blood from tail vein. Mice exceeded blood glucose levels 300 mg/dL after treatment were considered *diabetic* (HFD DM+).

### Diary food intake and mice weight measurements

To avoid mice distress interference because isolation and tail blood sampling, food and weight measurements were performed one week before ITT and GTT. Before measurements, mice were fasted for 12 hours overnight and then weighed for five consecutive days at the same hour. Food mass and water mass were also weighed at the same time for five consecutive days.

### Tissue extraction

Mice were submitted to fasting according to previously described protocol (45). Subsequently, mice were anesthetized with lethal doses of xylocaine and ketamine calculated by *Labinsane* App (43). Pancreas and fragments of dorsal skin (8.0 mm punch), epididymal and white adipose tissue were extracted, weighted and prepared for molecular analysis or histological staining.

### Glucose and insulin levels analysis

Plasma samples were obtained from fasted mice as here previously described (16). Whole blood samples collected in EDTA pre-coated tubes, followed by centrifugation (3500 RPM, 15 minutes, room temperature), and were stored at −80 °C. Serum glucose was determined by the glucose oxidase method, as previously described (17). Serum insulin was determined as previously described according to the manufacture’s instruction (Millipore #EZMI-13K)(16).

### Intraperitoneal glucose tolerance test

For Glucose tolerance test (GTT), mice were submitted to fasting protocol according previously described (45). After 6 h fasting, mice were anesthetized by an intraperitoneal injection of sodium amobarbital (15 mg/kg body weight). Basal glucose concentration was determined from collected tail blood. After collection of an unchallenged sample (time 0), a solution of 20% glucose (2.0 g/kg body weight) was administrated via intraperitoneal and blood from tail vein was collected after 15, 30, 60, 90 and 120 minutes. In both tests, blood glucose concentration was measured using handheld glucometer OptiumTM mini (Abbott Diabetes Care, Alameda, CA, USA) (46).

### Insulin tolerance test

For Insulin tolerance test (ITT) was performed as previously described (16). Briefly, mice were fasted and tail blood was collected for basal glucose evaluation. Fasted-mice received insulin (1.5 U/kg) by intraperitoneal injection, and blood samples were collected at 0, 5, 10, 15, and 20 min for glucose determination . The rate constant for glucose disappearance during insulin tolerance test (kITT) was calculated using the formula 0.693/t1/2. The glucose t1/2 was calculated from the slope of the least-square analysis of the plasma glucose concentrations during the linear decay phase (46). Blood glucose concentration was measured using handheld glucometer OptiumTM mini (Abbott Diabetes Care, Alameda, CA, USA).

### Histology

For the histological analyses, skin samples were processed and stained with hematoxylin and eosin as previously described (47). Briefly, skin and WAT samples were fixed by immersion in paraformaldehyde, processed in alcohol at 70, 80, 95, and 100%, followed by xylol and paraffin, embedded in paraffin blocks. Skin sections (5.0 μm) were placed on microscope slides pretreated with poly-l-lysine. We incubated sections with hematoxylin for 30 s, rinsed in distilled water, incubated for 30 s with eosin, rinsed again in distilled water, and dehydrated (47). Indirect white adipose tissue in skin sections were determined by the percentage of subcutaneus layer compared to the skin full thickness using H&E skin biopsies in five different animals in triplicate. Indirect hepatic steatosis was determined by the percentage of white ballooning formations in H&E liver biopsies in five different animals in triplicate (48, 49).

### Wound healing documentation and analysis

Full-thickness wounds were made as previously described (50, 51). Briefly, we used a sterile 6 mm dermal biopsy punch to perform two full-thickness wounds in the back skin of mice. The injury motion consisted of a single, quick perpendicular stroke accompanied by one rotatory movement to free the punch from the skin. The wound cavity was immediately dressed using a transparent bandage (Tegaderm) as previously described (52). For wound healing analyses, macroscopic evaluation of the healing process was evaluated as previously described (50, 51). Photographs were taken at 0, 4, 6, 8, 10 and 12 days after wounding using a wide camera 26 mm f/1.6 objective. The same distance (from the wound to the objective lens), top white fluorescence illumination, and operator were used on each occasion.. We calculate Wound Area as follows: Wound Area (percentage) = [(wound surface of day 0 - wound surface of day x)/wound surface of day 0] × 100. In each group, we use 16 to 20 photographs per day of analysis. Wound healing analysis were examined by ImageJ Software (1.52v) (51).

## AUTHOR CONTRIBUTIONS

CPJ: Conceptualization, Formal analysis, Methodology, Investigation, Writing - Original Draft, Writing - Review & Editing

GN: Conceptualization, Formal analysis, Methodology, Investigation JM: Conceptualization, Formal analysis, Methodology, Investigation TPP: Methodology and Investigation

RMB: Methodology and Investigation

LAV: Formal analysis, Writing - Review & Editing, Supervision WV: Formal analysis, Writing - Review & Editing, Supervision

EPA: Conceptualization, Methodology, Formal analysis, Resources, Writing - Original Draft, Writing - Review & Editing, Supervision

