## Supplementary figures and images for "A Diabetic Mice Model For Studying Skin Wound Healing"

### Supplementary 1

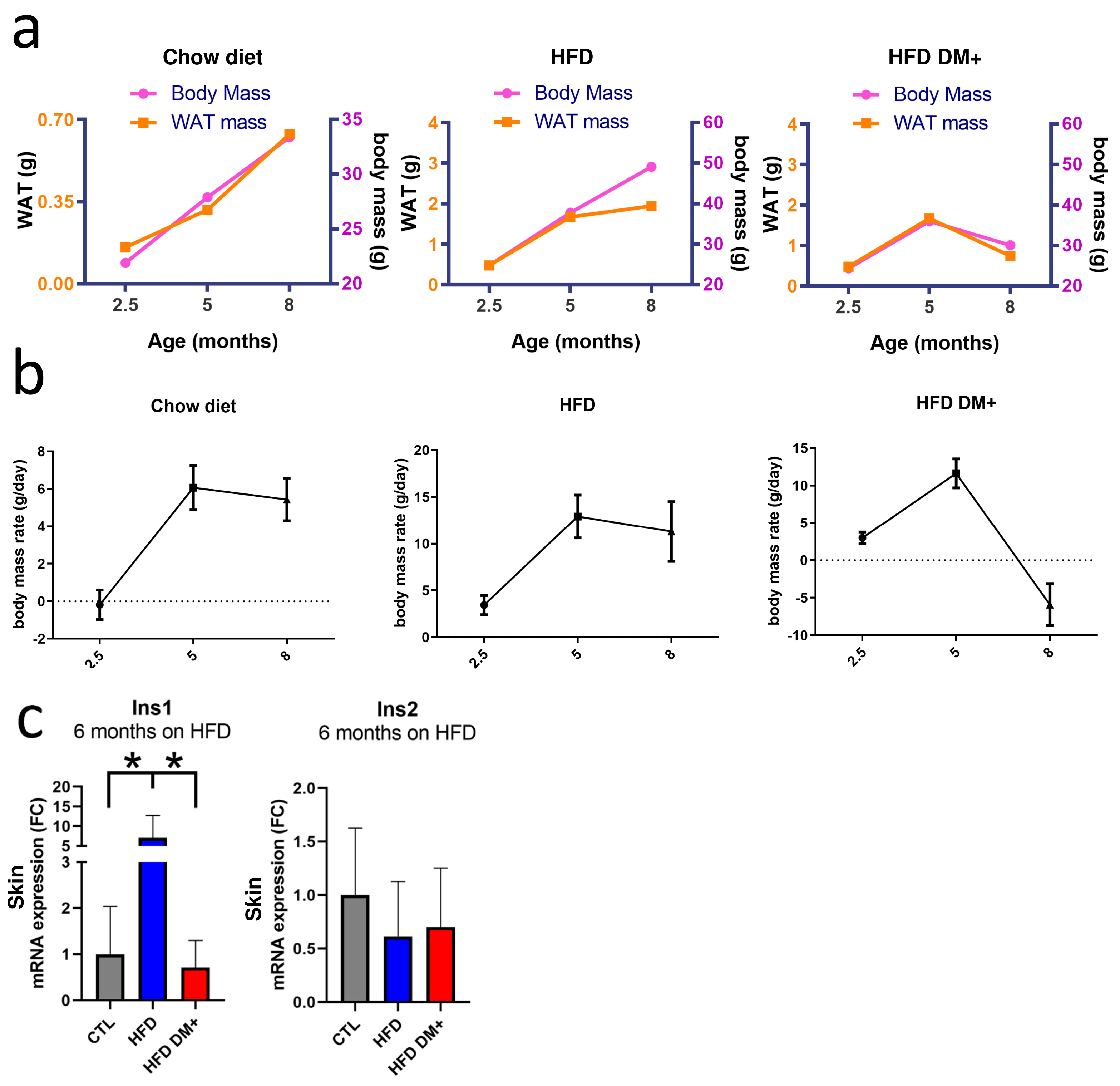
